# A human iPSC-based *in vitro* neural network formation assay to investigate neurodevelopmental toxicity of pesticides

**DOI:** 10.1101/2023.01.12.523741

**Authors:** Kristina Bartmann, Farina Bendt, Arif Dönmez, Daniel Haag, Eike Keßel, Stefan Masjosthusmann, Christopher Noel, Ji Wu, Peng Zhou, Ellen Fritsche

**Affiliations:** IUF - Leibniz Research Institute for Environmental Medicine, Duesseldorf, Germany; NeuCyte Inc., Mountain View, CA, USA; Medical Faculty, Heinrich-Heine-University, Duesseldorf, Germany; DNTOX GmbH, Duesseldorf, Germany

**Keywords:** developmental neurotoxicity, microelectrode arrays, electrical activity, human induced pluripotent stem cells, new approach methodologies

## Abstract

Proper brain development is based on the orchestration of key neurodevelopmental processes, including the formation and function of neural networks. If at least one key neurodevelopmental process is affected by a chemical, an adverse outcome is expected. To allow a higher testing throughput than the guideline animal experiments, a developmental neurotoxicity (DNT) *in vitro* testing battery (DNT IVB) has been set up that includes a variety of assays, which model several key neurodevelopmental processes. Gap analyses of the DNT IVB revealed the need of a human-based assay to assess neural network formation and function (NNF). Therefore, here we established the human NNF (hNNF) assay. A co-culture comprised of human-induced pluripotent stem cell (hiPSC)- derived excitatory and inhibitory neurons, as well as primary human astroglia, was differentiated for 35 days on micro-electrode arrays (MEA) and spontaneous electrical activity, together with cytotoxicity, was assessed on a weekly basis after washout of the compounds 24 h prior to measurements. In addition to the characterization of the test system, the assay was challenged with 28 compounds, mainly pesticides, identifying their DNT potential by evaluation of specific spike-, burst- and network parameters. This approach confirmed the suitability of the assay for screening environmental chemicals. Comparison of benchmark concentrations (BMC) with an NNF *in vitro* assay (rNNF) based on primary rat cortical cells, revealed differences in sensitivity. Together with the successful implementation of hNNF data into a postulated stressor-specific adverse outcome pathway (AOP) network associated with a plausible molecular initiating event for deltamethrin, this study suggests the hNNF assay as a useful complement to the current DNT IVB.

## 1 Introduction

The developing central nervous system is known to be more sensitive to exposure to toxic agents than the adult equivalent (Rodier, 1995). There is evidence that environmental chemicals contribute to neurodevelopmental disorders in children such as autism spectrum disorder, mental retardation, and cerebral palsy (National Research Council, 2000; Grandjean and Landrigan, 2006; Kuehn, 2010; Sagiv et al., 2010; Bennett et al., 2016). Pesticides belong to one compound class demonstrably associated with causing developmental neurotoxicity (DNT) (Bjørling-Poulsen et al., 2008). Today, only 35 of the 485 pesticides currently approved in the EU have been tested in DNT studies (Ockleford et al., 2018). The reason for this lack of testing, which generally expands to all chemicals (Goldman and Koduru, 2000; Crofton et al., 2012) lies in the current DNT *in vivo* testing guidelines: the OECD 426 (OECD, 2007) or EPA 870.6300 guideline (U.S. EPA, 1998). Their high resource intensity regarding time, money, and animals substantiate the limited throughput of these studies (Smirnova et al., 2014). Furthermore, high variability and low reproducibility of *in vivo* experiments, as well as species differences, increase the uncertainty of *in vivo* guideline studies for DNT testing (Tsuji and Crofton, 2012; Terron and Bennekou, 2018; Sachana et al., 2019; Paparella et al., 2020). In the last years, scientists from academia, industry, and regulatory authorities across the world agreed on the need for a standardized *in vitro* testing strategy, aiming for a cheaper and faster generation of additional data for DNT hazard assessment (EFSA, 2013; Crofton et al., 2014; Bal-Price et al., 2015, 2018; Fritsche, Crofton, Hernandez, Hougaard Bennekou, et al., 2017; Fritsche, Barenys, et al., 2018; Fritsche, Grandjean, et al., 2018). Following this consensus, a DNT *in vitro* battery (IVB) was compiled, which includes not one, but various DNT test methods, covering different neurodevelopmental processes, so-called key events (KEs), and developmental stages to approximate the complexity of human brain development (Fritsche, 2017; Fritsche, Crofton, Hernandez, Bennekou, et al., 2017; Bal-Price et al., 2018). Within this DNT IVB, neurodevelopment is described by *in vitro* assays covering the following KEs: human neural progenitor cell (hNPC) proliferation (Baumann et al., 2014, 2015; Harrill et al., 2018; Nimtz et al., 2019; Masjosthusmann et al., 2020; Koch et al., 2022) and apoptosis (Druwe et al., 2015; Harrill et al., 2018), cell migration (Baumann et al., 2015, 2016; Nyffeler et al., 2017; Schmuck et al., 2017; Masjosthusmann et al., 2020; Koch et al., 2022), hNPC-neuronal (Baumann et al., 2015; Schmuck et al., 2017; Masjosthusmann et al., 2020; Koch et al., 2022) and oligodendrocyte differentiation (Fritsche et al., 2015; Dach et al., 2017; Schmuck et al., 2017; Masjosthusmann et al., 2020; Klose et al., 2021; Koch et al., 2022), neurite outgrowth (<u>human</u>: Harrill et al., 2010, 2018; Krug et al., 2013; Hoelting et al., 2016; Masjosthusmann et al., 2020; Koch et al., 2022; <u>rat</u>: Harrill et al., 2013, 2018), as well as neuronal maturation and synaptogenesis (<u>rat</u>: Harrill et al., 2011, 2018).

Another crucial key neurodevelopmental process, also represented within the DNT IVB is the formation and function of neural networks, since the nervous system development requires functional networks consisting of different types of neurons and glial cells (Brown et al., 2016; Frank et al., 2017; Shafer, 2019). Furthermore, certain brain disorders, like autism spectrum disorder, Alzheimer’s and Parkinson’s disease, are associated with dysfunctional neural synchronization (Uhlhaas and Singer, 2006). Important tools to study electrophysiology of such neural networks are microelectrode arrays (MEA), which record extracellular local field potentials on multiple electrodes thus at different locations of the network and provide information on electrical activity, firing patterns, and synchronicity of the neural networks (Johnstone et al., 2010). So far, DNT *in vitro* testing for synaptogenesis and neuronal activity is mainly performed in assays based on rat primary cortical cells (Brown et al., 2016; Frank et al., 2017). The use of a human cell model to assess this endpoint has been identified as a gap in the current DNT IVB, precisely because the potential for species-specific features is still unknown (Crofton and Mundy, 2021).

The introduction of human-induced pluripotent stem cells (hiPSC) (Takahashi et al., 2007) has extensively advanced the field of biomedical sciences including testing for DNT. It has been proven that hiPSC-derived neural networks growing directly on MEAs exhibit spontaneous neuronal activity with organized spiking and bursting patterns (Odawara et al., 2016; Ishii et al., 2017; Nimtz et al., 2020; Tukker, Wijnolts, et al., 2020; Bartmann et al., 2021), which can be further modulated with known neurotoxicants and drugs (Odawara et al., 2018; Nimtz et al., 2020; Tukker, Bouwman, et al., 2020). The neural induction of hiPSCs towards functional neuronal cultures comes with many advantages, especially with regard to disease modelling but bears the issue of high variability between batches and cell lines. This variability is mostly due to the fact, that every single neural network differentiates into a variable number of neuronal subtypes. In addition, the generation of sufficiently active networks takes weeks to months (Hofrichter et al., 2017; Hyvärinen et al., 2019). The usage of commercially available hiPSC-derived neurons circumvents these problems with quality- and cell ratio-controlled, reproducible cells in large quantities (Little et al., 2019).

In this study, we present the establishment of a human neural network formation (hNNF) assay based on a commercially available kit, which consists of hiPSC-derived excitatory and inhibitory neurons and primary astroglia (SynFire, NeuCyte, USA). Pharmacological modulation confirmed the functionality of both neuronal subtypes and chronic treatment over 35 days revealed the ability of the cell model to detect alterations by bisindolylmaleimide I (Bis-I) through a known mode of action (MoA). Moreover, the assay was challenged with a test set of 28 substances and displayed compound-specific effects on network development.

## 2 Material & Methods

### 2.1 Compounds

In the present study, 28 substances were tested with various concerns regarding their DNT potential. Compounds are part of a training set used for the current DNT *in vitro* battery (Blum et al., 2022; Carstens et al., 2022). More information on the compound selection can be found in Masjosthusmann et al. (2020). As an assay negative control, Acetaminophen was included in the test set (assay negative control). The protein kinase C (PKC) inhibitor Bis-I was used as an assay positive control, together with bicuculline (BIC) and cyanquixaline (6-cyano-7-nitroquinoxaline-2,3-dione; CNQX) for acute pharmacological treatment of networks. Bis-I is known to decrease neurite outgrowth and firing/bursting rates of rat neural networks (Harrill et al., 2011; Robinette et al., 2011), whereas BIC and CNQX are GABAergic and glutamatergic receptor inhibitors, respectively. Compounds were dissolved in dimethyl sulfoxide (DMSO) or water to a stock concentration of 20 mM with exception of rotenone (100 mM), BIC (15 mM), and CNQX (30 mM). Applied concentrations ranged from 0.027 to 20 µM and 0.0004 to 0.3 µM for rotenone. Bis-I was applied at 5 µM, BIC at 3 µM, and CNQX at 30 µM. CAS registry numbers (CASRN), suppliers, and further information are collected in **Tab. S1**.

### 2.2 Cell Culture

SynFire glutamatergic neurons (Lot#000172 and 000131), SynFire GABAergic neurons (Lot#000172 and 000131) and SynFire astrocytes (Lot#13029-050 and 00190820; all from NeuCyte, USA) were thawed and cultured according to the manufacturer’s protocol. In short, all three cell types were thawed and resuspended in a defined ratio in supplemented seeding medium (NeuCyte), resulting in a co-culture consisting of induced neuronal (iN) and glial cells (iN:glia). Cells were seeded at a density of 270 × 10^3^ cells/well (140 × 10^3^ glutamatergic neurons, 60 × 10^3^ GABAergic neurons, 70 × 10^3^ astrocytes) on 48 well MEA plates (Axion M768-KAP-48) pre-coated with 0.1% polyethyleneimine and 20 µg/ml mouse laminin. This approach resulted in a cell-type ratio of 52% glutamatergic neurons, 22% GABAergic neurons and 26% astroglia. The seeding was performed in a 50 µl droplet (270 × 10^3^ cells/droplet) of supplemented seeding medium (NeuCyte) per well. After cells were allowed to adhere for 24 hours each well was filled with 250 µl supplemented short-term medium (NeuCyte). At days *in vitro* (DIV) 3 and 5 cells were fed by changing half of the medium with supplemented short-term medium. From DIV 7 onwards, medium was gradually changed to supplemented long-term medium (NeuCyte) by replacing half of the medium at each of DIV 7 and 10 until the medium was completely replaced at DIV 13 (washout). The short-term medium is supplemented with a substance that reduces the proliferation of astroglia. Following the same plating procedure and seeding density, 20 wells of a pre-coated 96-well flat bottom plate (Greiner) were prepared for weekly cytotoxicity assessments (see section 2.5).

### 2.3 Experimental Design

Following the first recording of spontaneous electrical network activity at DIV 7, cells were exposed to the respective test compound by changing half of the medium with supplemented long-term medium containing double-concentrated compound. Half medium changes with the compounds were conducted at DIV 10, 17, 24, and 31. The removed medium was used for cytotoxicity assessment using the CytoTox-ONE homogenous membrane integrity (LDH) assay (see 2.5). 24 hours before weekly recordings at DIV 7, 14, 21, 28, and 35 a washout with PBS was performed prior to replacing the medium with chemical-free supplemented long-term medium to minimize acute substance effects during MEA recordings. For this purpose, the medium was completely removed and replaced by adding 300 µl pre-warmed PBS. After incubation for 30 minutes (37°C, 5% CO_2_), PBS was replaced by freshly supplemented long-term medium. After recording, the medium was again replaced with long-term medium containing the test compound. Two compounds were tested per 48-well MEA plate including solvent and endpoint-specific controls. Each independent experiment (biological replicate) results from a different thawing procedure done on a different day and composes three technical replicates (replicate wells, see **Fig. S1**). In this study, we followed a two-step testing paradigm, where primarily each compound was tested twice independently. If the two independent experiments showed the same results, e.g no effect, no additional experiment was conducted. In case of conflicting outcomes, a third experiment was performed. The experimental setup is summarized in **Figure 1**.

**Figure 1:**
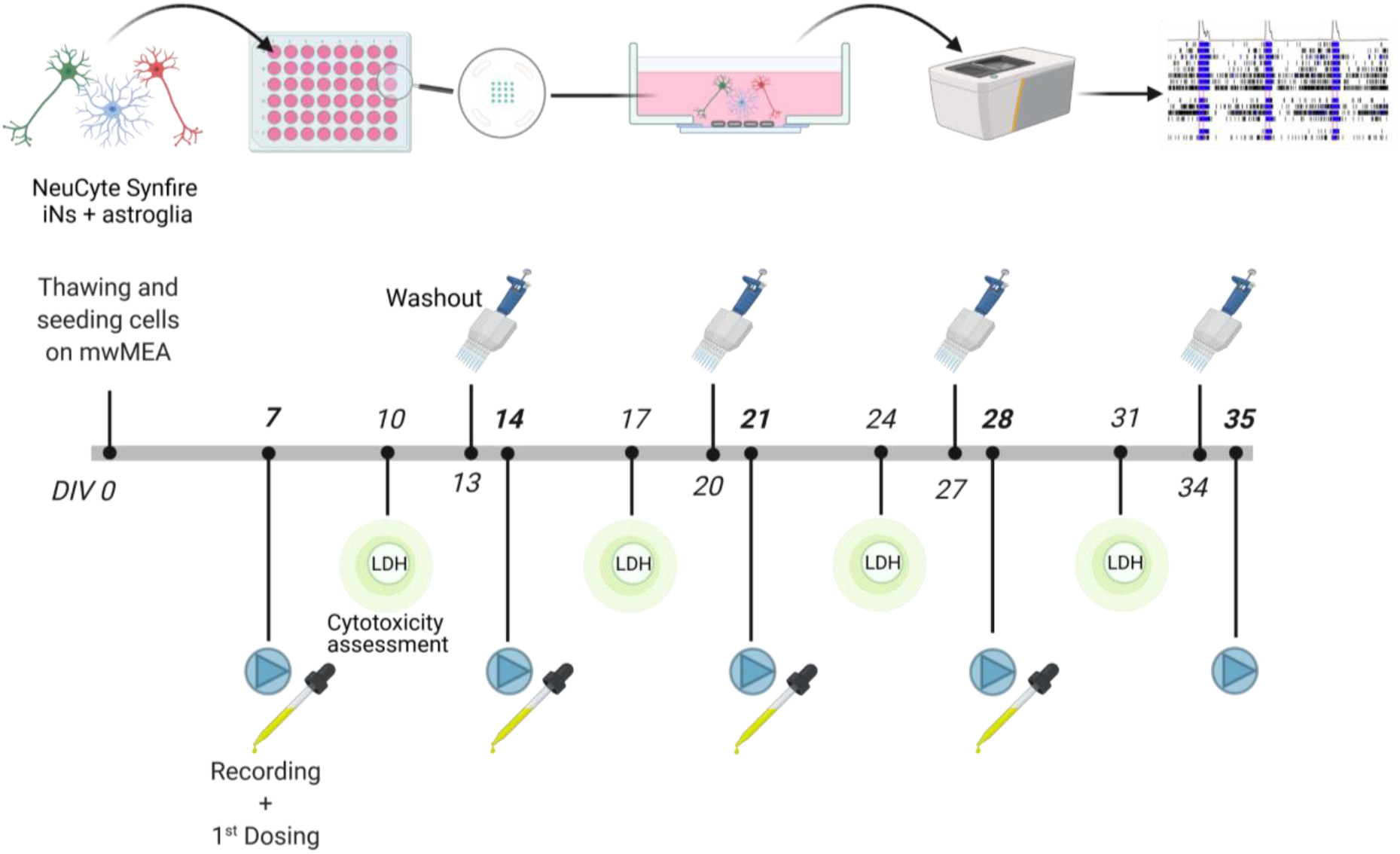
Experimental Setup of the human NNF assay. A co-culture of hiPSC-derived excitatory and inhibitory neurons and primary astroglia (NeuCyte, USA) was plated in a defined cell type ratio on 48-well MEA plates at DIV 0. Cultures were allowed to mature for 7 days before exposure to the test compounds. 24 hours before the weekly recording of spontaneous electrical network activity on DIV 7, 14, 21, 28, and 35 a washout of the respective compounds was performed. Additionally, cytotoxicity was assessed every week by the CytoTox-ONE (LDH) assay on DIV 10, 17, 24, and 31 three days after dosing.

### 2.4 MEA recording

Spontaneous electrical network activity on DIV 7, 14, 21, 28, and 35 was recorded with the Axion Maestro Pro system, a 768-channel amplifier, and the Axion Integrated Studio (AxIS) software version 1.5.3 or later (Axion Biosystems, Atlanta, USA). The recording procedure of the electrical network activity was composed of a 15 minutes equilibration period and two consecutive measurements of 15 minutes each, whereas only the last 15 minute recording was used for further analysis. For acute response measurements (CNQX, BIC) only the first 15 minutes recording was analysed. All recordings were conducted at 37°C and 5% CO_2_. The activity was measured using a gain of 1000 x and a sampling frequency of 12.5 kHz. A Butterworth band-pass filter was used (200 - 3000 Hz) prior to spike detection (threshold of 6x root mean square [RMS] noise on each electrode) via the AxIS adaptive spike detector. An active electrode was defined as ≥ 5 spikes/min.

### 2.5 Cytotoxicity assessment

Cytotoxicity was assessed every week and three days after re-dosing (at DIV 10, 17, 24, and 31) using the CytoTox-ONE Homogeneous Membrane Integrity assay according to the manufacturer’s instructions (CytoTox-ONE Homogeneous Membrane Integrity Assay; #G7891, Promega, Madison, United States). 50 µl medium from each well was removed, transferred to a 96-well plate (Sarstedt) and 50 µl CytoTox-ONE reagent was added. 30 minutes prior to the cytotoxicity assay, three wells of the lysis plate were treated with 10% Triton-X 100, and the supernatant was used as lysis positive control. As a background control, 50 µl of supplemented long-term medium was incubated with the same volume of CytoTox-ONE reagent. Following 2 hours of incubation at room temperature, the fluorescence was detected with a Tecan infinite M200 Pro reader (ex: 540 nm; em: 590 nm).

### 2.6 Data analyses

After recording with the Axion Integrated Studio (AxIS) software, recordings were re-recorded using the same software, resulting in .spk files. For single electrode burst detection, the Inter-Spike Interval (ISI) Threshold Algorithm was used with a maximum ISI of 100 ms with at least five spikes. Additionally, network bursts were detected using the Axion Neural Metric Tool and the Envelope algorithm with a threshold factor of 1.5, a minimum Inter-Burst Interval (IBI) of 100 ms, and 60% of active electrodes. The Synchrony Window was set to 20 ms. This resulted in 72 network parameters for five timepoints and seven concentrations. As the manual evaluation of all 72 parameters was not possible, an automated evaluation workflow that calculates the trapezoidal area under the curve (AUC) and benchmark concentrations (BMC) was set up. AUC was calculated as previously described by Brown and colleagues (Brown et al., 2016). Consecutively, spline interpolations with degree 1 polynomials for the data points given for conditions are made for each endpoint in each plate. If the response for an endpoint for the DIV 7 measurement was missing, e.g. because of absent bursting activity in an active well, it was set using random sampling throughout overall available first days-*in*-*vitro* responses for that endpoint on the same plate when at least 50% of these responses were available. In the final pre-processing step a common time duration, related to one DIV, with available responses in the resulting data is determined for each endpoint and experiment. Finally, the area under the resulting curves from these common durations was determined.

The pesticides data presented in this paper were derived from two to three individual experiments per compound as stated in **Tab. S1**. The data was normalized to the median solvent control and re-normalized to the starting point of a concentration-response curve generated with the R package drc as described below. For cytotoxicity data a different normalization was used: The normalized cytotoxicity response equalled the lysis control (LC) median minus the response of the respective concentration divided by the lysis control median minus the solvent control (SC) median 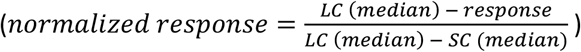. BMCs (BMC_50_ and BMC_50ind_) with their upper and lower confidence intervals (CI) were calculated based on the R package drc. Linear, sigmoidal, monotonic, and non-monotonic models were run with the concentration-response data of each endpoint, and Akaike’s information criteria were used to determine the best fit. Endpoints were classified as DNT-specific if CIs of the BMCs calculated for the DNT-specific endpoint did not overlap with the cytotoxicity endpoint. If the overlap exceeded 10%, the endpoint was classified as unspecific. Statistical significance was calculated using Graphpad Prism 8.2.1 and OneWay ANOVA with Dunnett’s post-hoc tests or two-tailed Student’s t-tests (p ≤ 0.05 was termed significant).

### 2.7 RNA-Seq

NeuCyte’s iPSC-derived glutamatergic, GABAergic induced neurons and human astrocytes were seeded to form iN:glia co-culture. Cells were harvested on DIV 7, 14, 21, 28, and 35, four biological replicates per time point. RNA-Seq was performed by Novogene (CA, USA). Total RNA was extracted by Qiagen’s RNA Extraction Kit (Qiagen, Germany). For library preparation, NEBNext® Ultra™ II RNA Library Prep Kit for Illumina® was used (New England Biolabs, MA, USA). For sequencing, NovaSeq 6000 was used, utilizing paired-end 150 bp read length. Downstream data analysis was performed using a combination of programs. Alignments were parsed using STAR program. Reads were aligned to the reference genome GRCh37 using STAR (v2.S). STAR counted number of reads per gene while mapping. The counts coincide with those produced by HTseq-count with default parameters. Then fragments per kilobase per million mapped reads (FPKM) of each gene was calculated based on the length of the gene and reads count mapped to this gene.

### 2.8 Immunostainings

SynFire iN:glia co-cultures were validated by immunostaining of markers including: Guinea pig anti-MAP2 (1:200, Synaptic Systems, 188 004), Rabbit anti-Synapsin1 (1:200, Synaptic Systems, 106 008), Rabbit anti-VGLUT2 (1:200, Synaptic Systems, 135 403), Rabbit anti-VGAT (1:200, Synaptic Systems, 131 011), Rabbit anti-GFAP (1:250, Abcam, ab4674). Secondary antibodies were conjugated to AlexaFluor647 (1:200, Invitrogen) and AlexaFluor488 (1:200, Invitrogen).

On DIV 35, the co-cultures were fixed with 4% paraformaldehyde (PFA; Sigma Aldrich) at room temperature for 15 min, washed three times with PBS (PAN Biotech), and then incubated overnight at 4°C with primary antibodies diluted in blocking solution [5% Goat Serum (Sigma Aldrich) + 0.2% Triton X-100 (Sigma Aldrich) in PBS]. On the second day, co-cultures were washed three times with PBS, and then incubated, with secondary antibodies diluted in blocking solution with 1% Hoechst 33258 (1:100, Merck). After three times washing with PBS, Fluorescence imaging was performed using an automated microscope system used for high content imaging (CellInsight CX7 LZR Platform, Thermo Fisher Scientific, Waltham, MA, USA).

## 3 Results

### 3.1 Characterization of an MEA-Based Assay for Network Formation

The present study describes the establishment and characterization of a human iPSC-based neural network formation assay. Furthermore, the assay was challenged with 27 pesticides and acetaminophen as a negative control. 270,000 cells of a defined cell type ratio (52% glutamatergic neurons, 22% GABAergic neurons, 26% astrocytes) were seeded as a monolayer culture on each MEA well containing 16 electrodes. This cell system is commercially available (NeuCyte, USA) and has been intensively characterized. **Figure 2** illustrates the characteristics of neural networks at different maturation time points (DIV 7-35) using transcriptome profiling (RNA-Seq), as well as immunocytochemical staining of DIV 35 networks. Stainings of differentiated co-cultures at DIV 35 (**Fig. 2A**) show a strong presence of MAP2-positive neurons and a lower amount of NeuN (*RBFOX3*), indicating a high maturation of the networks. In addition, the glial marker GFAP is strongly expressed. Furthermore, the co-cultures are positive for the pre-synaptic marker synapsin and exhibit the vesicular GABA transporter vGAT, as well as vGLUT, a vesicular glutamate transporter.

**Figure 2:**
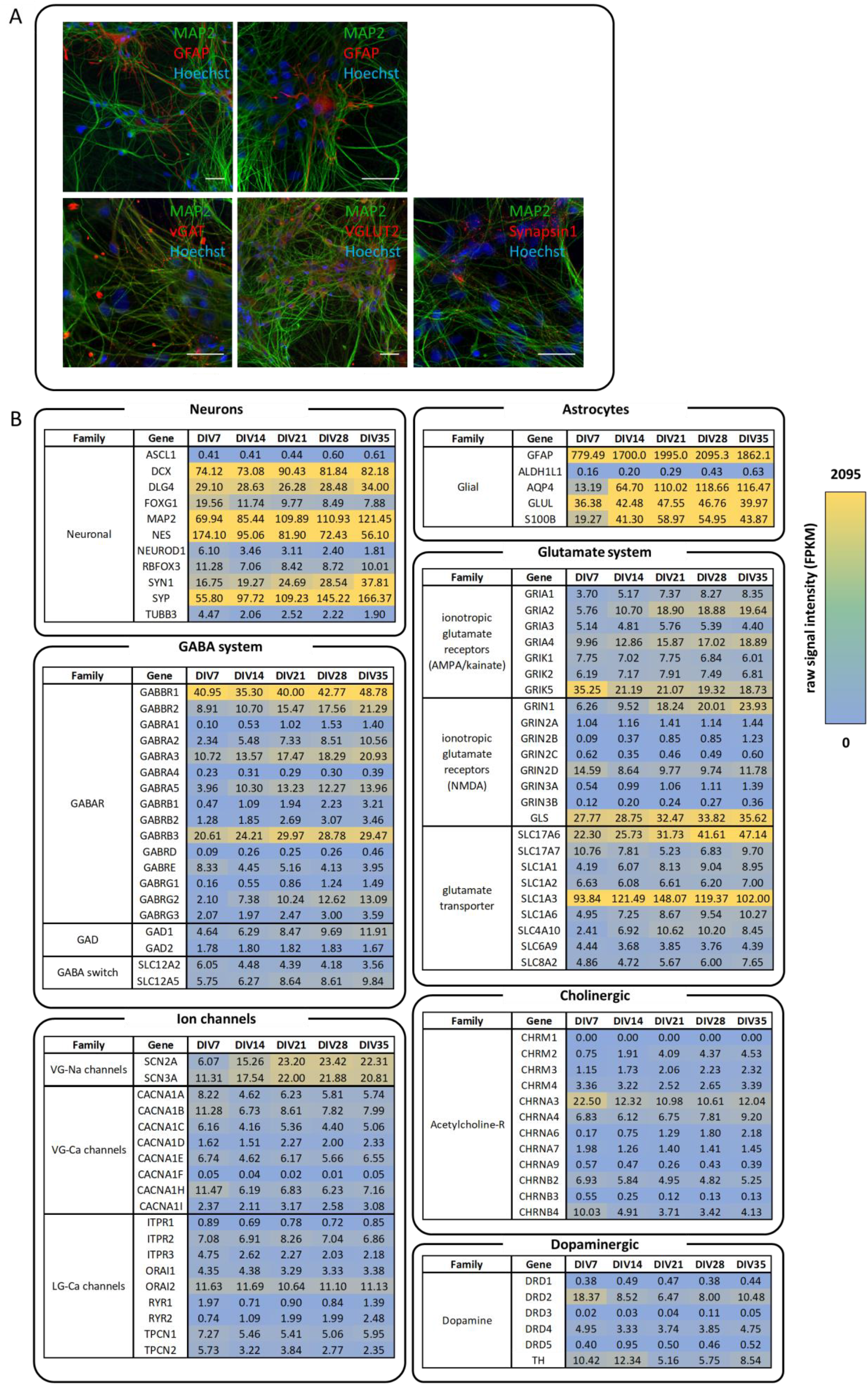
Immunocytochemical stainings and gene expression profiles of SynFire neuronal/glial co-cultures at different maturation time points. **(A)** Immunocytochemical stainings of different neuronal and glial markers of differentiated co-cultures at DIV 35. Nuclei were stained with Hoechst (blue) together with MAP2 (green) and either GFAP, Synapsin1, vGAT or vGlut, respectively (red). Scale bar = 50 µm. **(B)** RNA-Seq data was used to generate gene expression profiles on DIV 7, 14, 21, 28, and 35. Values are presented as fragments per kilobase per million mapped reads (FPKM). VG: voltage-gated; LG: ligand-gated; Ca: calcium; Na: sodium.

These characteristics are also confirmed by RNA sequencing of co-cultures at different time points (**Fig. 2B**). The proceeded maturation of neurons within the system is verified by a high expression of neuronal maturation markers, e.g. *MAP2* (Maccioni and Cambiazo, 1995), *DLG4* and *SYP* (Glantz et al., 2007), compared to genes, coding for immature neurons (e.g. *NEUROD1* (Seki, 2002)). Additionally, the high expression of *GFAP* and *AQP4*, compared to *S100B*, describes the mature glial system (Holst et al., 2019). Also, genes coding for GABA and AMPA receptors and glutamate transporters showed expression at DIV 7 with gradations within their subtypes (e.g. *SLC1A2* vs. *SLC1A3*). In addition, voltage- (VG) and ligand-gated (LG) ion channels are enriched in the culture. In comparison to VG- and LG-calcium channels, VG-sodium channels are higher expressed. Moreover, decent expression levels were detected for transcripts coding for dopaminergic, cholinergic (nicotinic and muscarinic), and NMDA receptors (NMDAR). Especially *CHRNA3* (Karlin, 2002) and *DRD2* (Missale et al., 1998) expressions are enhanced at DIV 7 and decrease with increasing maturation of the networks. The higher expression of *SLC12A5* (*KCC2*) compared to *SLC12A2* (*NKCC1*) at DIV 14 gives an indication that the cells are in a stage after the GABA switch, which marks the shift from pre-mature excitatory to mature inhibitory GABAergic neurons (Leonzino et al., 2016).

Taken together, these gene expression data show that the neural networks develop over time and express a broad variety of genes related to neuronal and glial function as a prerequisite for neural network function. Important tools to study electrophysiology of such networks are MEAs. MEA recordings provide high content data based on the recording of extracellular action potentials, so-called spikes, which are the basic unit of activity of a neural network. During the development of a network, spikes can group into bursts and also synchronize their activity resulting in network bursts. To confirm the contribution of functional GABAergic and glutamatergic neurons in the development of the networks, we performed an acute pharmacological modulation of neural networks on DIV 21. Therefore, the neuronal subtypes included in the cell model were challenged with the two receptor antagonists BIC and CNQX. BIC is a well-studied GABA_A_ receptor antagonist, which primarily competitively inhibits binding of GABA to its receptors. As a consequence, BIC reduces GABA-activated conductance by reducing channel opening times, as well as the opening frequency, resulting in an increased firing rate (Macdonald et al., 1989; Johnston, 2013; Mack et al., 2014). In comparison, CNQX primarily antagonizes AMPA-type glutamate receptors and thereby suppresses spontaneous excitatory glutamatergic synaptic activity (Neuman et al., 1988; Odawara et al., 2016). After baseline recording at DIV 21, cells were exposed to the respective modulator and the electrical activity was measured subsequently. Exposure to 3 µM BIC increased general electrical activity, especially synchronous bursting (**Fig. 3A**, pink boxes), whereas 30 µM CNQX led to a loss of organized activity, as illustrated by representative 100-second spike raster plots (**Fig. 3A, B**). For additional quality control of the test method, acute treatments with the two described modulators were included in every assay run. Violin plots for each compound and four different network parameters (**Fig. 3C-F**) illustrate the distribution of these data and the median of 8-9 independent experiments (median of 3 technical replicates each) reveals the effect of both receptor antagonists. 3 µM BIC significantly enhanced the mean firing rate and burst duration. Consistent with the spike raster plots, BIC doubled the percentage of overall spikes contributing to network bursts from 40% to 80% (“Network burst percentage”, **Fig. 3E**) and increased the synchronicity (“AUNCC”, **Fig. 3F**). In contrast, CNQX inhibited the overall activity and organization of the networks. Especially burst duration and synchronicity were impaired with a low degree of variation (**Fig. 3H, J**).

**Figure 3:**
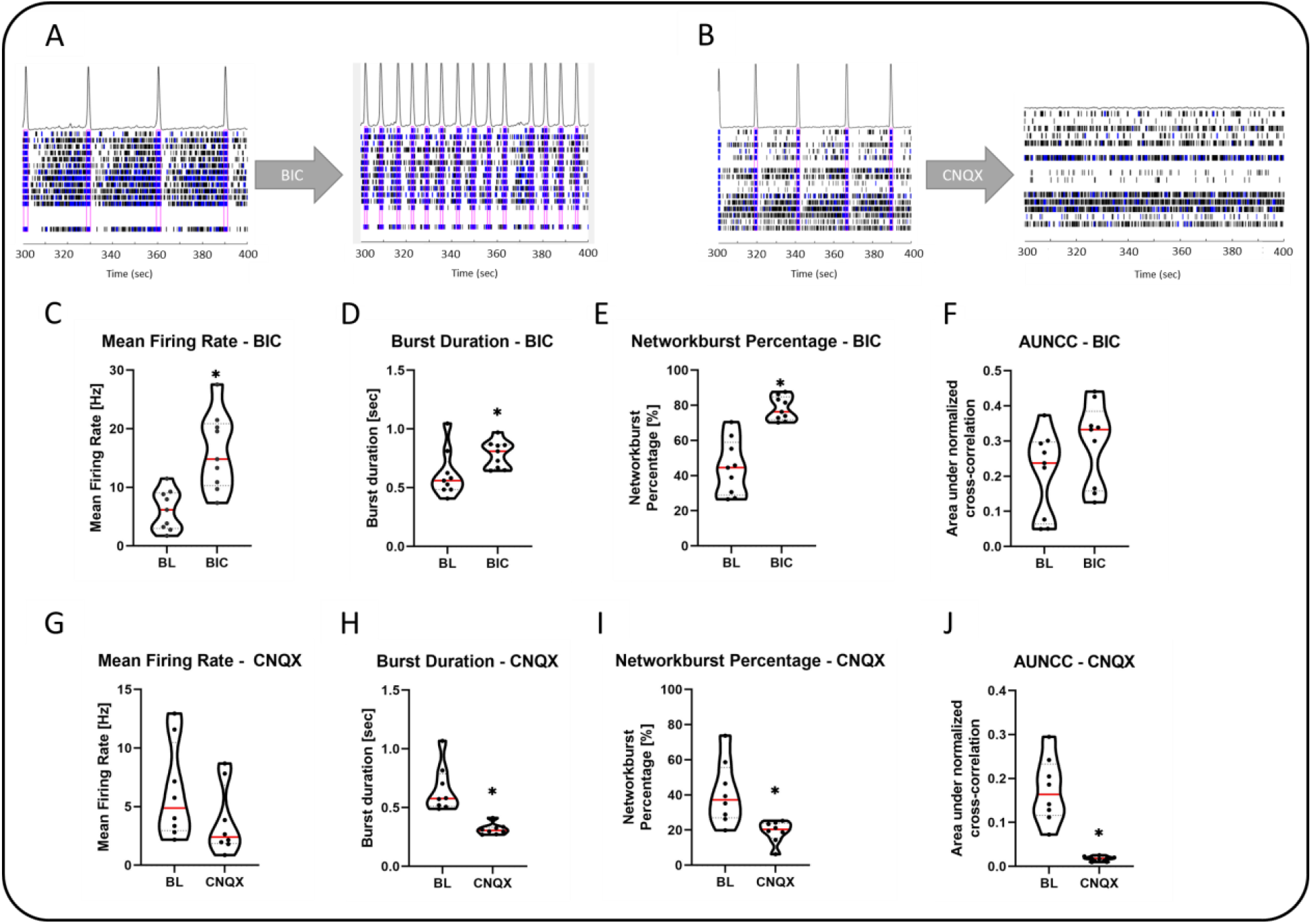
Acute pharmacological modulation of DIV 21 neural network activity. Untreated DIV 21 neural networks (baseline, BL) were exposed to 3 µM bicuculline (BIC) and 30 µM cyanquixaline (6-cyano-7-nitroquinoxaline-2,3-dione; CNQX), respectively. **(A, B)** 100-second spike raster plots reveal the effects of 3 µM BIC and 30 µM CNQX on DIV 21 neural networks. Spikes are represented as black bars, bursts as blue bars. Pink boxes indicate network bursts. **(C-F)** 3 µM BIC enhanced neural network activity as indicated by the increase of different network parameters compared to the baseline. **(G-J)** 30 µM CNQX decreased neural network activity through different network parameters. Data are represented as violin plot distribution of 9 **(C-F)** or 8 **(G-J)** independent experiments. Dotted lines represent quartiles of distribution and red bars the median. Statistical significance was calculated using two-tailed Student’s t-tests. A p-value below 0.05 was termed significant. *significant compared to the respective BL.

As already indicated by gene expression data (**Fig. 2**), electrophysiological measurements over time confirm maturation of the neural networks by displaying specific firing patterns, like organized spiking and synchronicity at later points of differentiation (**Fig. 3A**). Representative spike raster plots illustrate these features over the 35-day development of the networks (**Fig. 4A**). At DIV 7 spikes (black bars) are the sole form of activity, whereas at DIV 14 bursts (blue bars) start to form on single electrodes. Along with the increase in bursting activity and the emergence of a synchronous network starting on DIV21, the number of spikes between network bursts decreases at DIV 28 and 35. This transition in network development is reflected by the evaluation of different network parameters of untreated solvent control wells (0.1% DMSO) from all plates contributing to this study (n=41 plates and 123 wells). **Figure 4B-E** shows the distribution of different parameters between experiments (each data point reflects the median of 3 wells of each independent experiment) and indicates the variation of the hNNF assay. On DIV 7 and 14, the mean firing rate is notably below 5 Hz, but raises to 10 Hz, with its peak at DIV 28 (**Fig. 4B**). The same trend can be observed for the duration of bursts between DIV 7 and 35 (**Fig. 4C**). Nevertheless, some parameters, like the network burst percentage or the area under normalized cross-correlation, which describes the synchronicity of the network, continuously increase and find their peak at DIV 35 (**Fig. 4D, E**). Because the highly organized activity of the network is rarely observed at DIV 7 and 14 (**Fig. 4E**), network burst parameters were only considered from DIV 21 to 35 for the following evaluations.

**Figure 4:**
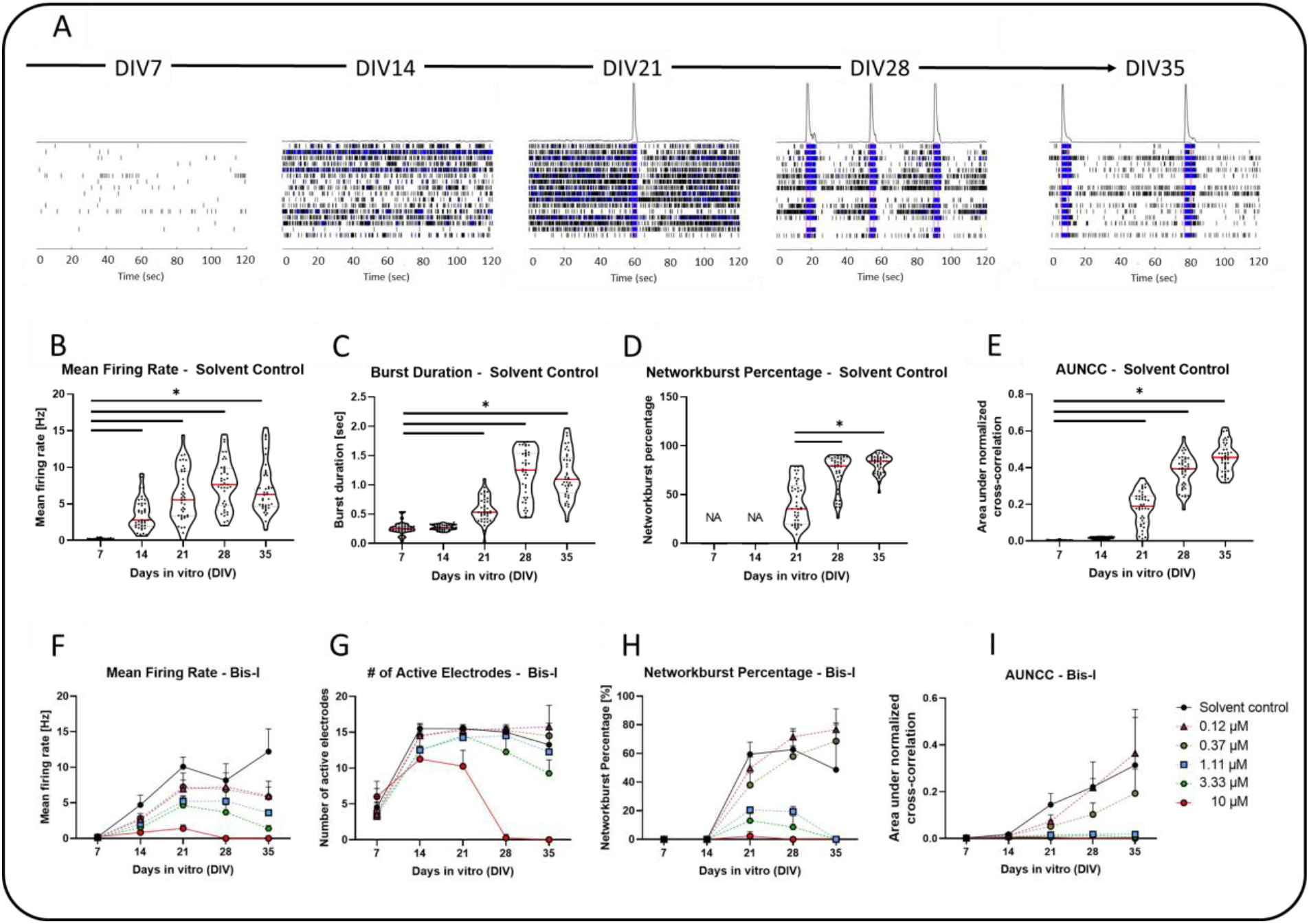
Neural network development on 48-well microelectrode arrays (MEA) and its inhibition by Bisindolylmaleimide I (Bis-I). **(A)** Representative 120 second spike raster plots referring to DIV 7 to 35. Spikes are represented as black bars, bursts as blue bars. Pink boxes indicate network bursts. **(B-E)** Dot plots showing the distribution of untreated (solvent control; 0.1% DMSO) network activities from DIV 7 to 35 over four network parameters (mean firing rate, burst duration, network burst percentage, and area under normalized cross-correlation (AUNCC)). Single dots represent the median of three replicates of each experiment. The red bar defines the median overall plates (n=41). **(F-I)** Starting at DIV 7 networks were treated with increasing concentrations (0.12; 0.37; 1.11; 3.33 and 10 µM) of the PKC inhibitor Bis-I. Different network parameters (mean firing rate, number of active electrodes, network burst percentage, area under normalized cross-correlation (AUNCC)) reflect the impairment of Bis-I on neural network development over 35 days of differentiation. Data are shown as mean ± SD of four wells. Statistical significance was calculated using one-way ANOVA. A p-value below 0.05 was termed significant. **\***significant compared to the earliest DIV.

To show that neural network development within the hNNF assay can be altered by a specific mechanism, DIV 7 networks were exposed to different concentrations of the protein kinase C inhibitor Bis-I, following the exposure scheme described in **Figure 1**. Bis-I inhibits neurite outgrowth in PC-12 cells (Das et al., 2004), rat cortical neurons, and human iPSC-derived neurons (Druwe et al., 2016). Furthermore, *in vitro* MEA experiments showed decreased firing and bursting rates of rat neural networks after Bis-I exposure (Robinette et al., 2011). We observed that network activity was affected by Bis-I in a concentration-dependent manner, as illustrated by representative network parameters in **Figure 4F-I**. Untreated controls showed a mean firing rate of more than 10 Hz on DIV 35, whereas Bis-I interfered with the formation of a functional network starting at low concentrations of 0.12 µM resulting in about 5 Hz (**Fig. 4F**). Exposure to 10 µM Bis-I resulted in a fully muted network at DIV 28 and DIV 35 observable in every of the four displayed network parameters. Not only the general activity was affected, but also the network burst percentage was reduced to 20% at DIV 21 and 28 by 1.11 µM Bis-I and culminate in 0% at DIV 35 (**Fig. 4H**). This impairment in network bursting is also reflected in the area under normalized cross-correlation (AUNCC; **Fig. 4I**). After this initial proof-of-concept, Bis-I was introduced as an endpoint-specific positive control for the hNNF test method. For this purpose, each experimental run in the compound screening contained control wells, treated with 5 µM Bis-I. The data of 13 independent experiments (39 wells) showed that Bis-I reliably and significantly inhibits different network parameters, e.g. burst duration, network burst percentage and network synchronicity (AUNCC, **Fig. S2**). The overall effect of Bis-I on the mean firing rate was not significant, which may be explained by the higher variability of this parameter, as explained in the next section (see **Fig. S2**).

### 3.2 Selection of Parameters to Evaluate

MEA recordings generated in this study result in 72 network parameters, which are predominantly correlated and can be grouped into spike-, burst- and network-related parameters. A plethora of these define the same characteristic of the network (e.g., “mean firing rate” and “weighted mean firing rate”) or use a different statistical method to describe the parameter (e.g. “inter-burst interval - Avg” vs “inter-burst interval (median) - Avg”). The evaluation of one 48-well MEA plate during a time course of 35 days results in over 17,000 data points, which enormously exacerbates the processing of data and interpretation of possible compound effects. To reduce the number of data points and only concentrate on the most informative and at the same time robust parameters, we analyzed the variability of all parameters across all wells treated with the lowest concentration of each test compound. We therefore calculated the inter-experimental variability for each parameter, by merging the endpoint responses of the lowest concentrations of independent experiments relative to the solvent control and normalized these. Afterwards, we derived the inter-experimental variability as the coefficient of variation (COV) of these collections of data points for each compound. The higher the COV, the greater is the dispersion. Additionally, we included parameters that were previously described in the literature (Brown et al., 2016; Frank et al., 2017; Kosnik et al., 2020) and which represent a broad variety of network development. We then selected a final set of 14 network parameters, that covers the three categories “General activity”, “Bursting activity” and “Connectivity” of the neural networks and shows a COV between 6 and 29 **(<u>Table 1</u>)**.

**Table 1:** 14 Parameters from MEA recordings and their respective inter-experimental variability as coefficient of variation (COV).

| Category | Parameter | Definition | Inter-experimental COV |
| --- | --- | --- | --- |
| General Activity | Mean Firing Rate | Total number of spikes divided by the duration of the analysis [Hz] | 28.45 |
|  | Number of Active Electrodes | Number of electrodes with activity > 5 spikes / minute | 9.97 |
|  | Number of Bursting Electrodes | Total number of electrodes within the well with bursts/minute greater than the burst electrode criterion (min# of spikes: 5; max ISI: 100 ms) | 11.14 |
| Bursting Activity | Burst Duration | Average time from the first spike to last spike in a single-electrode burst | 11.24 |
|  | Number of Spikes per Burst | Average number of spikes in a single-electrode burst | 25.17 |
|  | Mean ISI within Burst | Mean inter-spike interval, time between spikes, for spikes in a single-electrode burst | 5.94 |
|  | Inter-Burst Interval | Average time between the start of single-electrode bursts | 19.18 |
|  | Burst Frequency | Total number of single-electrode bursts divided by the duration of the analysis [Hz] | 28.60 |
|  | Burst Percentage | The number of spikes in single-electrode bursts divided by the total number of spikes, multiplied by 100 | 14.67 |
| Connectivity | Network burst Frequency | Total number of network bursts divided by the duration of the analysis [Hz] | 20.83 |
|  | Network burst Duration | Average time from the first spike to last spike in a network burst | 11.42 |
|  | Network burst Percentage | The number of spikes in network bursts divided by the total number of spikes, multiplied by 100 | 7.08 |
|  | Number of Spikes per Network burst | Average number of spikes in a network burst | 29.01 |
|  | Area Under Normalized Cross-Correlation (AUNCC) | Area under the well-wide pooled inter-electrode cross-correlation normalized to the auto-correlations | 14.73 |

### 3.3 Concentration-Dependent Effects of Pesticides on Network Activity

After having set up the hNNF assay by setting up a defined treatment scheme and standard operation procedure, establishing an endpoint-specific control and evaluating variability over wells and plates, we next applied the hNNF assay for screening of 28 chemicals. The set consists of 27 pesticides and acetaminophen as a negative control compound. To identify concentration-dependent effects of substances that impaired neural network formation, cells were weekly exposed from DIV 7 to 35, including respective washout steps 24 hours before each recording. As an example, deltamethrin and β-cyfluthrin reduced the mean firing rate **(Fig. 5A)** and area under normalized cross-correlation **(Fig. 5B)** of neural networks, respectively, in a time- and concentration-dependent manner without inducing cytotoxicity. Acibenzolar-S-methyl, on the other hand, did not affect the number of active electrodes **(Fig. 5C)**. These time-concentration relationships can be translated into concentration-response curves as illustrated in **Figure 5D-F**. For each concentration, the trapezoidal area under the curve was calculated to include all five time points in one single value per concentration. This approach, as adapted from Brown et al. (2016) and Shafer et al. (2019) simplifies the comparison of effects over different days of neural network development. In the next step, the AUC information and resulting concentration-response curves were used to estimate BMCs with upper and lower confidence limits for each compound and parameter. For estimation of the BMCs a bench mark response (BMR) of 50% (BMR_50_) was selected as this best reflects the variability of the most variable parameters **(see Tab. 1)**.

**Figure 5:**
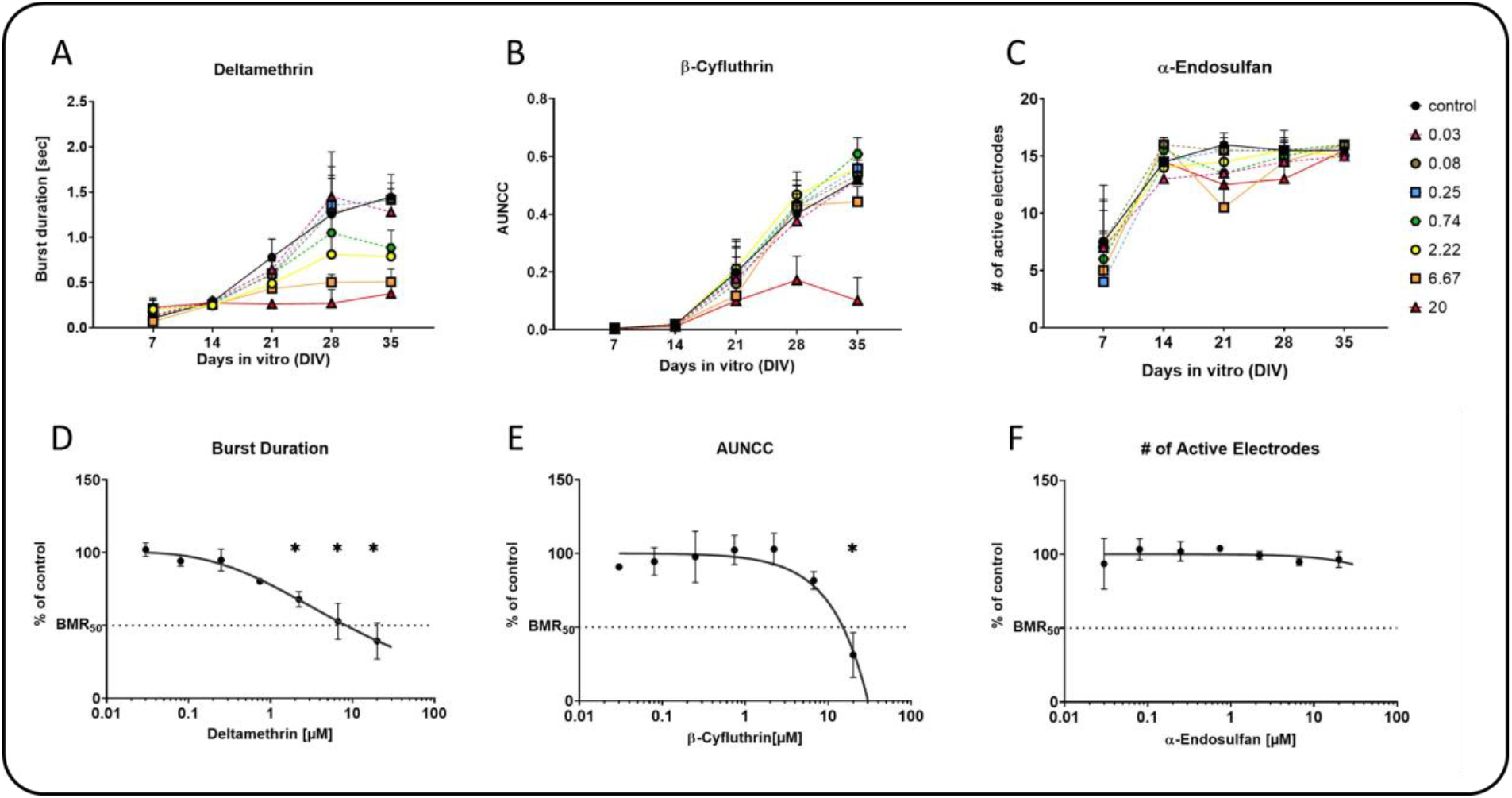
Example data for area under the curve (AUC) summary of time- and concentration-dependent MEA readouts. **(A-C)** Starting at DIV 7, neural networks were treated with increasing concentrations of deltamethrin **(A)**, β-cyfluthrin **(B)**, and α-endosulfan **(C)**, and exemplary network parameters were evaluated. Time- and concentration-dependent data are shown as the mean of three **(A, B)** or two **(C)** independent experiments ± SEM **(A, B)** or SD **(C)**. Area under curve values were computed for these data and plotted in a concentration-dependent relationship **(D-F)**. Compounds did not induce cytotoxicity (data not shown). Data are represented as the mean of three **(D, E)** or two **(F)** independent experiments ± SEM **(D, E)** or ± SD **(F)**. Replicates within one expriment are summarize by median. Curve fitting was conducted as described in section 2.6. Statistical significance was calculated using one-way ANOVA. A p-value below 0.05 was termed significant. **\***significant compared to the respective solvent control.

**Figure 6** summarizes the concentration-dependent effects of the 28 tested compounds on neural network development that produced a 50% change (reduction or induction) from the curves starting point. Five of the 14 network parameters were not affected by any compound, including number of active electrodes and mean ISI within bursts. Furthermore, 17 compounds were tested negative, e.g. acetaminophen, chlorpyrifos, and its derivate chlorpyrifos-methyl (data not shown). 11 of the 27 pesticides are considered positive, for which at least one network parameter had to be affected, without cytotoxic effects at any administered concentration. β-cyfluthrin, β-cypermethrin, deltamethrin, penthiopyrad, and rotenone evoked effects in more than two parameters, whereas the other six pesticides affected one or two parameters. Network burst frequency is characterized as the most sensitive parameter with six hits, of which five represent inductive effects. Furthermore, spirodiclofen and penthiopyrad additively increased the burst frequency. Predominantly, the observed effects are all in a similar range between 9 and 20 µM. In contrast, deltamethrin and metaflumizone influenced different network parameters below 9 µM. Rotenone is denoted as the most potent compound, specifically reducing seven parameters in a concentration range between 0.15 and 0.24 µM without causing cytotoxicity.

**Figure 6:**
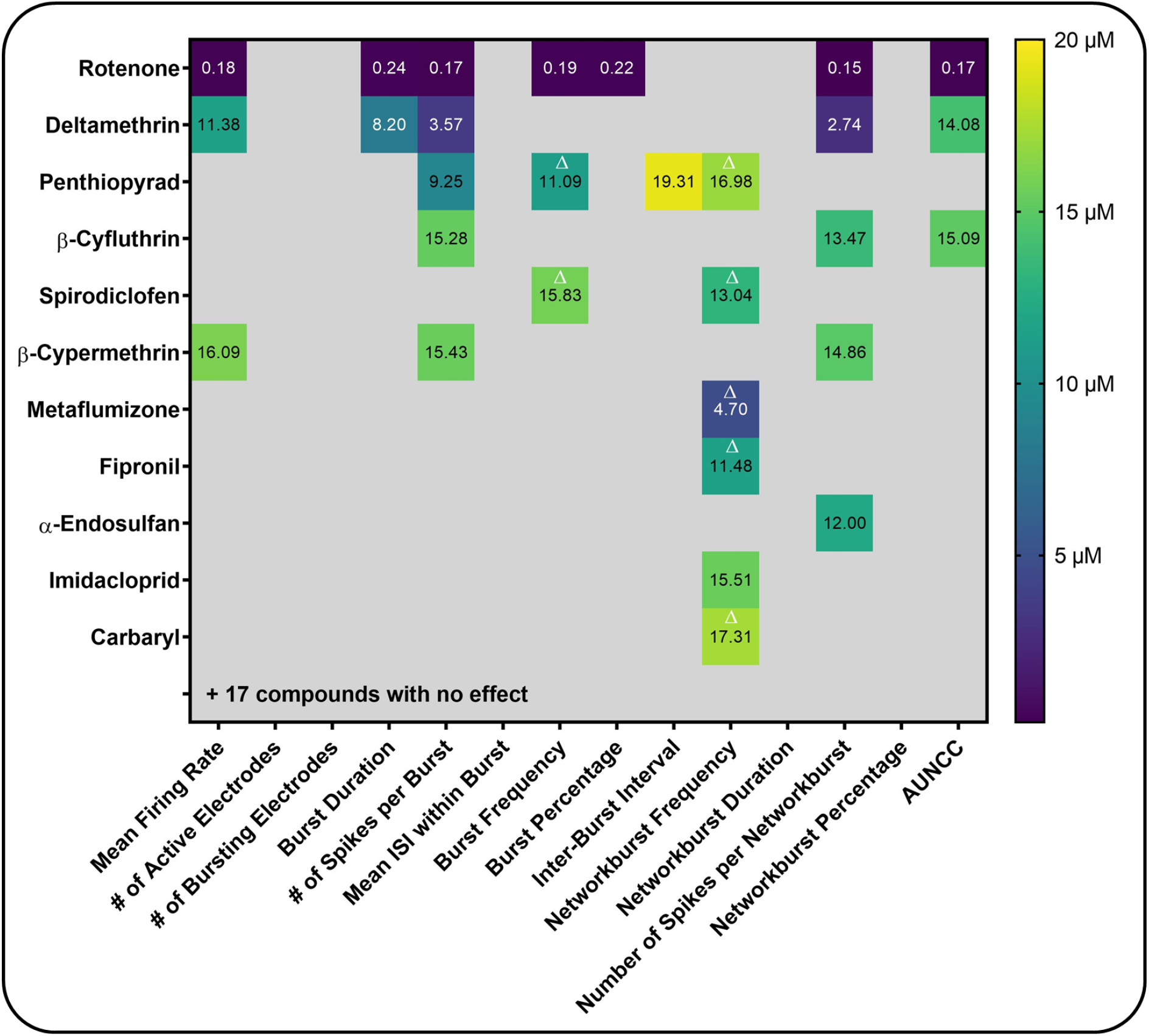
Summary of BMCs across 14 network parameters of the hNNF assay. No cytotoxicity was observed. 17 compounds had no effect (acetaminophen, acetamiprid, acibenzolar-s-methyl, aldicarb, chlorpyrifos, chlorpyrifos-methyl, clothianidin, diazinon, dimethoate, dinotefuran, disulfoton, etofenprox, flufenacet, methamidophos, thiacloprid, thiamethoxam, triallate; data not shown). Δ Induced effects. Numbers are given in µM. No value assumes BMCs > 20 µM (> 0.3 µM for rotenone). Confidence intervals are listed in **Tab. S2**. AUNCC: area under normalized cross-correlation.

## 4 Discussion

In the last years, scientists from academia, industry, and regulatory authorities across the world agreed on the need for a standardized *in vitro* testing strategy, aiming for a cheaper and faster generation of additional data for DNT hazard assessment (EFSA, 2013; Crofton et al., 2014; Bal-Price et al., 2015, 2018; Fritsche, Crofton, Hernandez, Hougaard Bennekou, et al., 2017; Fritsche, Barenys, et al., 2018; Fritsche, Grandjean, et al., 2018). Therefore, a DNT IVB was compiled, including various test methods covering different KEs of neurodevelopment, including the formation and function of neural networks (Fritsche, 2017; Fritsche, Crofton, Hernandez, Bennekou, et al., 2017; Bal-Price et al., 2018). One of the identified gaps of the DNT IVB is the assessment of network formation and function in a human-based cell model (Crofton and Mundy, 2021). This is why we established the human neural network formation assay (hNNF), which consists of hiPSC-derived excitatory and inhibitory neurons and primary astroglia (SynFire, NeuCyte, USA). By pharmacological modulation, the functionality of neuronal subtypes and the ability of the cell model to detect alterations by a MoA were confirmed. Furthermore, the assay was challenged with a test set of 28 substances and revealed compound-specific effects of different pesticides on network development.

### Assay establishment

Under most circumstances, newly developed methods for DNT testing are restricted by their ability to test large numbers of chemicals (Coecke et al., 2007; Crofton et al., 2011). To tackle this issue, Crofton et al. provided a set of 15 principles, which should enhance the amenability of higher throughput screening (Crofton et al., 2011). The establishment of the hNNF assay in this study was realized by considering these principles, which are described in more detail in the following (P1-15). During early brain development, neurons start to mature and build connections via synapses (Okado et al., 1979; Zhang and Poo, 2001). Neural network formation and function is therefore a key aspect of neurodevelopment (P1 “Key Event of Neurodevelopment”). By measuring extracellular local field potentials on MEAs, network formation and function can be assessed, thus providing information on electrical activity, firing patterns, and synchronicity of the neural networks (P2 “Endpoint Measurement”). By calculating the AUC for each concentration and normalizing the values to the respective solvent control, the hNNF assay can reflect alterations of network activity in both directions (increase and decrease; P3 “Dynamic Range”). Furthermore, Crofton et al. emphasize the importance of parametric controls, meaning parameters of the assay that evoke predictable changes in the endpoint (P4 “Parametric Controls). One aspect that was confirmed within the presented study is the increase in electrical activity and synchronicity of the networks with increasing culture time (**Fig. 3**). Furthermore, Saavedra and colleagues showed that an excitatory:inhibitory (ex:inh) ratio of 70:30 using SynFire neurons exhibits the steadiest spiking increase and coverage of electrodes over differentiation time, compared to other ex:inh ratios (Saavedra et al., 2021). Principle 5 (“Response Characterization”) highlights the relevance of a precise effect characterization, based on the degree of variability in the assay. As recommended by the EFSA Scientific Committee, the BMC approach was applied, to derive a reference point or point of departure (Hardy et al., 2017), whereby the BMR should be defined as an effect size that is higher than the general variability of the measured endpoint. Based on the inter-experimental standard deviation (1.5*COV), which was calculated for every parameter presented in this study (**Tab. 1**), we defined the BMR_50_ (reduction) and BMR_50ind_ (induction) as the degree of change that, if exceeded, results in a positive response (hit). Furthermore, Crofton and colleagues state, that the concentration range and the resulting concentration-response holds a very significant role in terms of comparison of sensitivity between different endpoints. For this study, we selected a concentration range that has already been chosen in other *in vitro* DNT assays and has elicited little to no cytotoxicity (Frank et al., 2017; Masjosthusmann et al., 2020; Blum et al., 2022). To discriminate specific from unspecific effects, we assessed the cytotoxicity of each compound on a weekly level during the 35-day culture period (P7 “Endpoint Selectivity”). Another crucial requirement for assay development is the selection of endpoint-specific controls, altering the endpoint by known MoA, both negatively and positively. In the present study, Bis-I, a PKC inhibitor was selected as an endpoint-specific control (P8 “Endpoint-selective controls). In primary rat cortical cells the inhibition of PKC blocked the local astrocytic contact and thus the facilitation of excitatory synaptogenesis throughout the neuron (Hama et al., 2004). In particular, in MEA experiments, Bis-I decreased the firing and bursting rates of rat neural networks *in vitro* (Robinette et al., 2011). Within the hNNF assay, Bis-I reliably reduced network parameters e.g. burst duration, network burst percentage and network synchronicity (AUNCC). Due to the enhanced variability of the mean firing rate, no significant effect of Bis-I was observed over all conducted experimental runs. Nevertheless, Bis-I is an appropriate endpoint-specific control for assessing neural network activity *in vitro*. As a negative control compound, acetaminophen showed no effect on network activity. A plethora of studies confirmed the use of acetaminophen as an apt negative control for DNT *in vitro* testing (Radio et al., 2008; Stern et al., 2014; Brown et al., 2016; Masjosthusmann et al., 2020). Nevertheless, a recently published study by Martin and colleagues categorizes acetaminophen as an unfavourable negative control for DNT assays (Martin et al., 2022). This classification was made based on both clinical and preclinical reports, which indicate potential effects on the developing brain. Furthermore, the authors state the usage of low concentrations of acetaminophen (≤ 100 μM) in recent DNT studies as a possible reason for the absence of compound-dependent effects. In comparison, chemicals like L-ascorbic acid, dinotefuran and metformin were listed as ‘favourable’ and should be included in a compound set for later determination of the assay’s general performance (Martin et al., 2022). Following the recommendations of Crofton and colleagues, a training set of chemicals should be designed and assayed (P9 “Training Set Chemicals”), after demonstrating that the test method has the aforementioned characteristics. Chemicals should be included that produce a reliable effect on the endpoint in focus and chemicals that do not, which allows both specificity and sensitivity of the assay to be determined (P11 “Specificity and Sensitivity”). The hNNF assay was established and used within a research project with a focus on pesticides. In consideration of the high cost involved in performing substances screening in the assay, it was not possible to distinguish between training and testing set of chemicals during the establishment process. Instead, we selected pesticides that have different DNT potentials, according to several *in vivo* and *in vitro* studies (P10 “Testing Set Chemicals”; Masjosthusmann et al. 2020). Pyrethroids, for example, are linked to epidemiological studies that report neurodevelopmental disorders during childhood after pyrethroid pesticides exposure (Oulhote and Bouchard, 2013; Xue et al., 2013; Pitzer et al., 2021). Especially deltamethrin is a thoroughly studied type II pyrethroid, for which animal studies reported long-term effects on the brain (summarized in Pitzer et al., 2021), which was also observed *in vitro* (Shafer et al., 2008; Masjosthusmann et al., 2020). In contrast, the neonicotinoid dinotefuran was described as DNT negative *in vivo* (Sheets et al., 2016) and also recommended as a negative tool compound for alternative DNT test methods (Aschner et al., 2017). In the future the hNNF assay will also be challenged with more chemicals that are well-described DNT positive and negative compounds to assess specificity and sensitivity of the assay and to enhance the readiness of the test method (Bal-Price et al. 2018). Currently, the academic setup of the hNNF assay allows parallel testing of 12 compounds (n=1) within the 35-day experimental period. However, it is possible to increase the throughput by increasing the plate size format from 48- to 96-well or by introducing automation. Furthermore, duplicating the experimental set-up by using multiple MEA recording devices or by shifting the recording days in a periodic manner, would also increase the throughput (P12 “HighThroughput”). Principle 13 (“Documentation”) highlights the importance of documenting the test method in detail to allow an easy transfer and implementation across laboratories. A detailed standardized protocol exists and is currently being transferred to a laboratory of the U.S. EPA. After initial establishment of the assay in the collaborating laboratory, the hNNF assay will be challenged with a set of test substances to inform about the robustness and inter-laboratory transferability of the assay. This point is also emphasized in principle 14 (“Transferability”), along with the guaranteed availability of the cells used, which can be purchased commercially by other researchers. Currently, the generated data is not being shared through an open access databased (P15 “Data Sharing”), but the authors aim to include hNNF data in a future ToxCast™ (http://www.epa.gov/ncct/toxcast) release. In summary, the hNNF assay fulfils the majority of the principles proposed by Crofton et al. (2011) for the establishment of *in vitro* DNT assays for substance screening. Currently, the low number of tested chemicals defines the lack of readiness of the assay (Phase I Readiness Score B, Phase II Readiness Score C; Bal-Price et al., 2018) and the improvements required for the assay to be ready will be tackled in the future by testing known DNT positive substances (Aschner et al., 2017).

### The hNNF assay compared to its rat counterpart

The hNNF assay was established to model neural network formation and function in a human-based cell model and to become a valuable addition to the current DNT IVB, which comprises 17 different test methods, able to measure changes in key neurodevelopmental processes (Masjosthusmann et al., 2020; Crofton and Mundy, 2021). Neural network formation and function is currently modelled in an assay based on rat primary cortical cells (rNNF), assessing the developmental effects of chemicals over 12 days of differentiation (Brown et al., 2016). It is important to mention that we aligned the hNNF assay with the parameter set of the rNNF assay and thus both assays provide comparable parameters of network development (e.g. number of active electrodes or burst duration) to reduce uncertainty. **Table 2** juxtaposes the results obtained in this study with rNNF data (Frank et al., 2017). Comparing the BMC_50_ values of the most sensitive endpoint (MSE) between the hNNF and rNNF assay, it becomes clear that the observed positive hits differ in sensitivity across all substances. The rNNF seems to be more sensitive as it detects effects on network activity even at lower concentrations of the tested substances, e.g. Deltamethrin (BMC_50_ MSE hNNF: 2.74 µM; BMC_50_ MSE rNNF: 0.5 µM). Aldicarb and chlorpyrifos were negative in the hNNF, but altered network formation in the rNNF assay. Acetaminophen was identified as a negative in both assays.

**Table 2:** Comparison of BMC_50_ values of the most sensitive endpoint (MSE) for chemicals (same CASRN) tested in the hNNF (this study) and rNNF assay (Frank et al., 2017). ↑ indicates an inductive effect (BMC_50ind_).

|  | Acetaminophen | Aldicarb | Carbaryl | Chlorpyrifos | Deltamethrin | Fipronil | Imidacloprid |
| --- | --- | --- | --- | --- | --- | --- | --- |
| **BMC<sub>50</sub> MSE hNNF** | no hit | no hit | 17.31↑ | no hit | 2.74 | 11.48 ↑ | 15.51 |
| **BMC<sub>50</sub> MSE rNNF** | no hit | 0.66 | 0.08 | 1.4 | 0.5 | 1.33 | 9.99 |

The hNNF and rNNF assay are referred to as complementary assays because they measure similar endpoints, i.e. several MEA parameters, but differ with regards to species (rat vs. human), presence of fetal bovine serum (FBS) and assay technology (beginning and wash-out of compounds before MEA recordings). Therefore, differences in data obtained within these assays are not necessarily evidential of a false detection (Crofton and Mundy, 2021). These differences may be justified by several distinctions in the assay setup, i.e. species differences and exposure schemes. The hNNF and rNNF assays are based on the same basic cell types, namely neurons and astrocytes, but derived from different species (hNNF: human iPSC-derived neurons and primary astroglia; rNNF: rat primary neocortical cells). It is widely accepted that the predictability of non-human-based assays for human health is limited by species differences (Leist and Hartung, 2013). Also primary neural progenitor cells (NPC) derived from rats (PND5), are more sensitive towards exposure with DNT compounds compared to time-matched primary human NPCs *in vitro* (Baumann et al., 2016). These two systems differ not only in their sensitivity but also with regard to their molecular equipment notwithstanding similar cellular functions (e.g. NPC migration and differentiation; Masjosthusmann et al., 2018). Recently, the co-culture system applied in this study was used for comparing acute effects of neurotoxic compounds on network activity to rodent cultures (rNNF) and revealed a considerable delay in human iPSC-derived neuronal and glial co-culture compared to rat cortical cultures (Saavedra et al., 2021). In general, the developing rat brain exhibits some crucial differences from human brain development *in vivo*, such as the absence of gyrification, which adds complexity to the human brain (Dubois et al., 2008). Furthermore it has been demonstrated, that embryonic day (E) 18 and E21 during rat brain development match with week 8-9 and week 15-16 after fertilization in human embryo, when looking at neurogenesis (Bayer et al., 1993). The faster maturation of rodent cells compared to human cells *in vitro* was also suggested by Masjosthusmann et al. (2018).

As the compound set presented in **Table 2** is rather small and focussed on pesticides, we cannot draw general conclusions about species-specific sensitivity of the rNNF and hNNF assays. As suggested by Bauman et al. (2016), testing of additional compounds with known MoA required to infer more general species-specific sensitivity. Nevertheless, our data highlights the importance of considering species specificities when comparing screening results.

Besides the species, experimental procedures differ between the human and rat NNF assays and may also lead to differences in assay sensitivity. Two major exposure differences are crucial. First, the timepoint when the compound is administered differs between hNNF and rNNF assay. Rat cortical cultures are exposed to the compound two hours after seeding the cell on MEAs, whereas the first day of dosing in hNNF experiments is DIV 7, when first single spike activities are visible. The respective networks are at different stages of development at this time, i.e. in contrast to human cells, rat cortical cells are barely established in the culture dish. Not yet established cells shortly after the plating process may be more susceptible to substance exposure than networks that have already been able to differentiate for a week in chemical-free medium. In the rNNF assay, early processes like neurite initiation and outgrowth, as well as glial proliferation are potentially disrupted within the first 24 hours (Harrill et al., 2011; Frank et al., 2017), whereas these processes can proceed undisturbed during the first seven days of differentiation and contribute to network development in the hNNF assay. Both assays thus depict different stages of neural network development and hence include different windows of neurodevelopmental processes. In addition, the hNNF culture medium is supplemented with FBS, which contains a variety of plasma proteins, peptides, and growth factors, whereas the rNNF medium is not. Recent studies have shown that FBS in cell culture medium attenuates the toxicity caused by specific compounds due to their high affinity for binding to proteins, which then results in higher BMC values (Zhang et al., 2016, 2020). The influence of FBS in the human cultures with regards to network effects needs to be studied by assessment of *in vitro* kinetics in the future (Kramer et al., 2015). Secondly, the washout of the respective compound 24 hours prior to the recording is a unique feature of the hNNF assay and aims at minimizing acute substance effects during MEA recordings. There is evidence that specific substances directly target synaptic receptors and acutely affect brain function. For example, the NMDAR is a prime target of the heavy metal lead, leading to the inhibition of glutamatergic synapse activity (Toscano and Guilarte, 2005). To diminish the measurement of these acute effects and to only assess the effects of substances on neural network development, we introduced washout steps into the experimental procedure of the hNNF. In comparison to the rNNF results (**Tab. 2**) it is possible that presence of compounds during MEA measurements contributes to higher sensitivity of the rat versus the human NNF assay.

All of the aforementioned variations in assay setup and biology, either alone or in combination, can explain the discrepancies in sensitivity between the two test methods. In the future, exposure schemes of the two NNF assays should be harmonized in order to understand the true nature of species differences concerning neural network formation. This might substantially help extrapolating from rat *in vivo* studies to humans using the parallelogram approach (Baumann et al. 2016).

### Use of hNNF data on deltamethrin for the development of a putative AOP

In 2021 the EFSA developed an IATA case study with the goal of including all available *in vivo* and *in vitro* data, among others the data generated within the DNT IVB for DNT hazard identification for the Type II pyrethroid insecticide deltamethrin (Crofton and Mundy, 2021; Hernández-Jerez et al., 2021). Epidemiological studies revealed associations between childhood exposure to pyrethroids like deltamethrin and neurodevelopmental disorders, e.g. attention deficit hyperactivity disorder or autism spectrum disorder (Oulhote and Bouchard, 2013; Shelton et al., 2014; Wagner-Schuman et al., 2015). As previously shown, deltamethrin negatively influenced 5 of 14 parameters describing network function with “Number of spikes per network burst” as the most sensitive endpoint within the hNNF assay (BMC_50_ 2.7 µM). Here, interference with voltage-gated sodium channels (VGSC) is the most commonly known MoA for pyrethroid insecticides like deltamethrin (Tapia et al., 2020), therefore representing one of two molecular initiating (MIE) events within the stressor-specific AOP network (**Fig. 7**). This MIE is followed by key events 1-6 and 9, describing different cellular responses, like the disruption of sodium channel gate kinetics, leading to disruption of action potential and in the end cumulate in an impaired behavioural function (adverse outcome). KE4 describes the alteration of neural network function as shown also by data assessed in the rNNF (BMC_50_ 0.5 µM; **Tab.2**) and hNNF assay. The 5-fold higher BMC of the hNNF assay compared to the rNNF assay might be explained by the different exposure paradigm and/or the different species as discussed above in more detail. Furthermore, potential mechanisms or processes that are disrupted by a chemical agent can be revealed and used for the development of adverse outcome pathways (AOP) and also set a new focus for more hypothesis-driven *in-vivo* studies (Hernández-Jerez et al., 2021). The postulated stressor-based AOP network (Fig. 7) is currently not included in the OECD AOP Wiki, but the EFSA Panel on Plant Protection Products and their residues recommends the submission to the OECD program to further support a regulatory uptake. This case study and the inclusion of hNNF data on deltamethrin exposure showed the applicability of the hNNF assay for hazard identification and characterization, consistent with the other assays of the DNT IVB.

**Figure 7:**
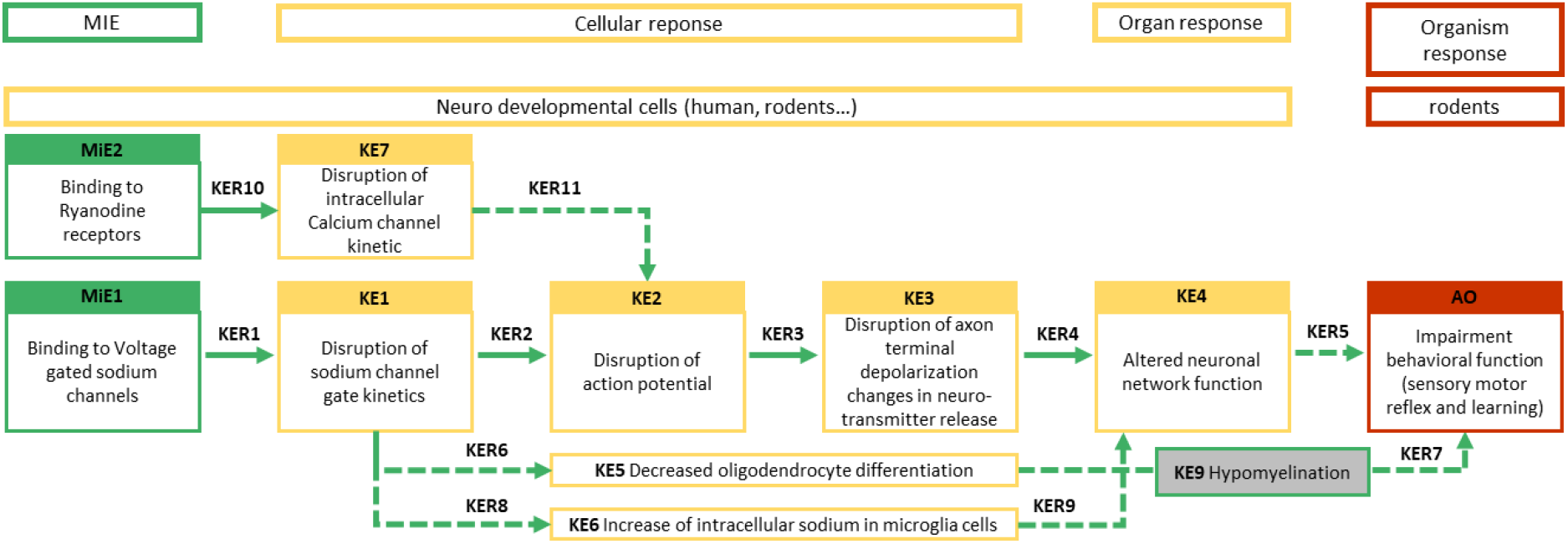
AOP network on deltamethrin postulated by the EFSA Panel on Plant Protection Products and their Residues. Non-adjacent key events for which the biological reasonability and/or empirical evidence is less assured, are marked by dashed lines. MIE: molecular iniating event; KE: Key event; KER: key event relationship; AO: adverse outcome. Adapted from Hernández-Jerez et al., 2021.

The presented study provides insight into the establishment of a novel new approach method, assessing alterations on neural network formation and function, using an hiPSC-derived co-culture of neurons and primary astroglia. The cell model comprises a broad variety of genes expressed exclusively in neurons and astrocytes as a prerequisite for neural network function after the GABA switch. For example, together with the rNNF assay, it is capable of representing NMDAR expression and assessing any MoA involving NMDAR, which distinguishes these NNF assays from other assays of the DNT IVB (Masjosthusmann et al., 2020). Compared to other studies using excitatory and inhibitory iNs the hNNF performs similarly with regard to e.g. firing rates and number of active electrodes. For example, Saavedra and colleagues cultured the excitatory and inhibitory iNS, which resulted in 14 to 16 active electrodes after 37 days and a mean firing rate between 1 and 2 Hz after 21 days. Furthermore, the described test system was successfully used to assess acute neuroactive effects of chemicals (Saavedra et al., 2021; Tukker et al., 2020). Nevertheless, beside astrocytes, also other cell types play an important role in development and function of neural networks, such as oligodendrocytes, responsible for myelination (Doretto et al., 2011) or microglia, which, among other things, phagocyte weak synapses and regulate neurogenesis (Miyamoto et al., 2016; Paolicelli and Ferretti, 2017). To represent all processes appropriately, the hNNF assay has to be complemented with the missing cell types. Still, a proactive establishment of the assay provided already a medium readiness of the assay for use in regulatory screening approaches and the testing of 28 substances revealed the suitability of the assay for screening environmental chemicals, like pesticides. In the future, the throughput of the hNNF assay as well as its robustness and specificity will be increased by testing additional substances, thereby enlarging the chemical space, to present a suitable addition to the current DNT IVB and close one of the identified gaps regarding network formation and function.

## Availability of data

The dataset generated during and/or analyzed during the current study is available from the corresponding author upon reasonable request.

## Supplementary Material

**Table S1:**
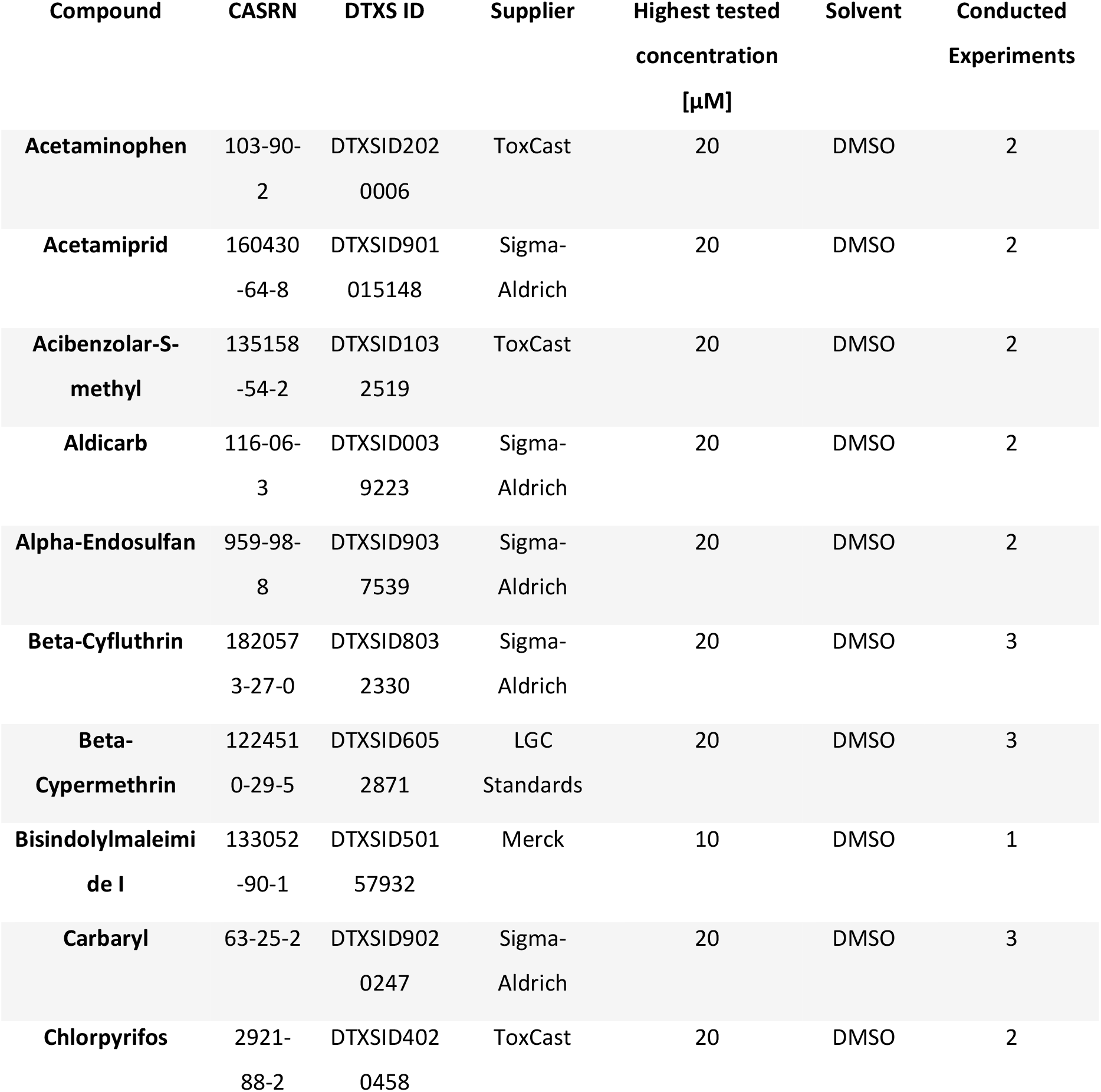
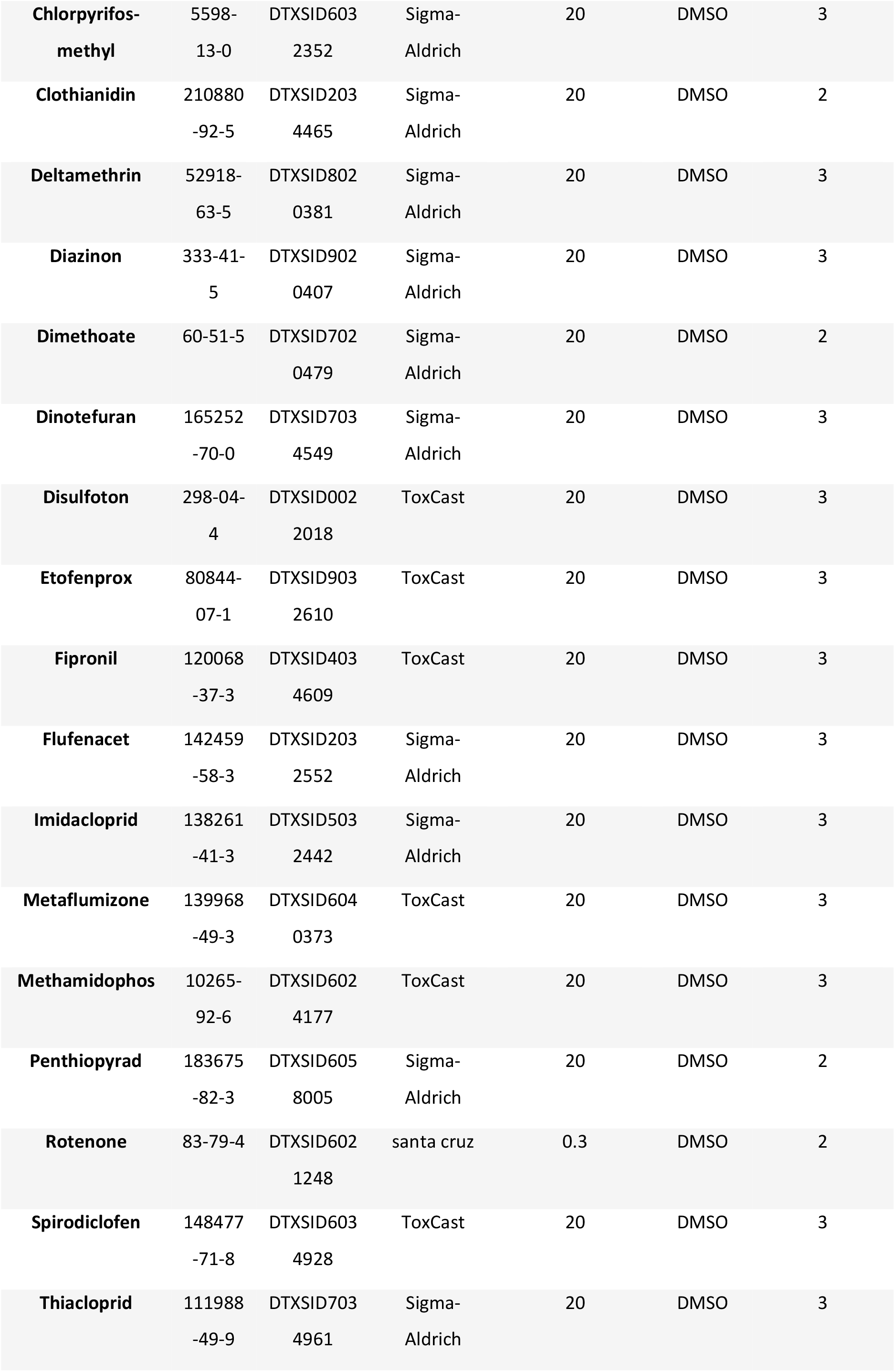
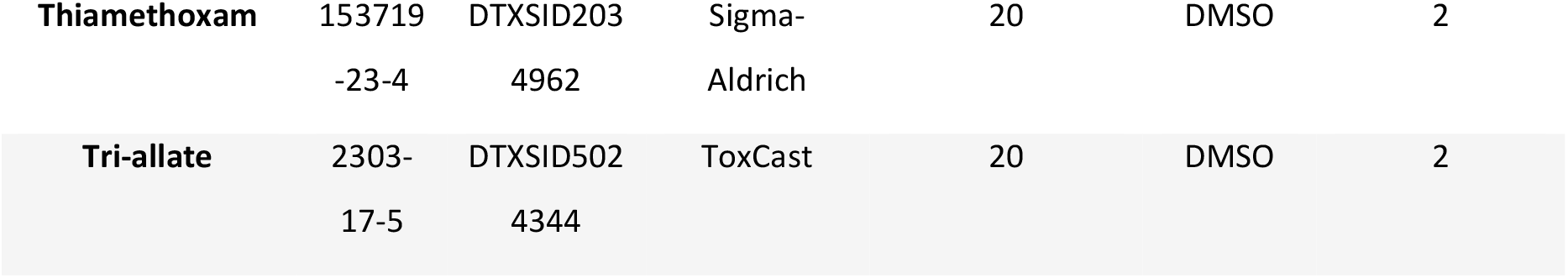
Chemical Compounds used for toxicity testing on MEAs. CASRN: CAS Registry Number. DTXS ID: DSSTox substance identifier.

**Figure S1:**
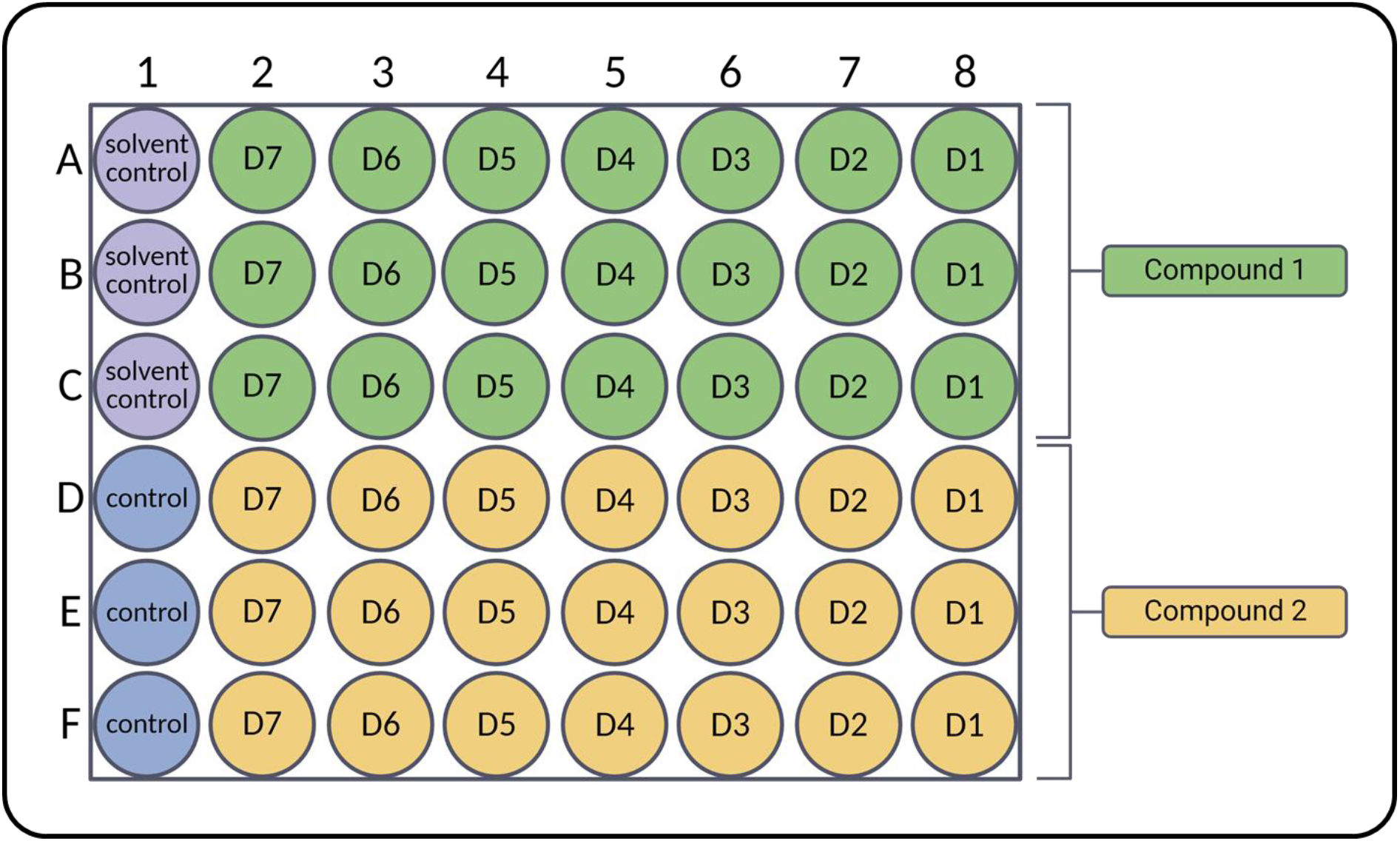
Exposure scheme of the hNNF assay. Solvent control: 0.1% DMSO. D7-1: Dilution 7 (lowest concentration) – dilution 1 (highest concentration). Controls include Bis-I exposure (5 µM) during the whole assay duration, as well as acute treatment with Bicuculline and CNQX at DIV21. For each experimental run at least three wells for each control were used.

**Figure S2:**
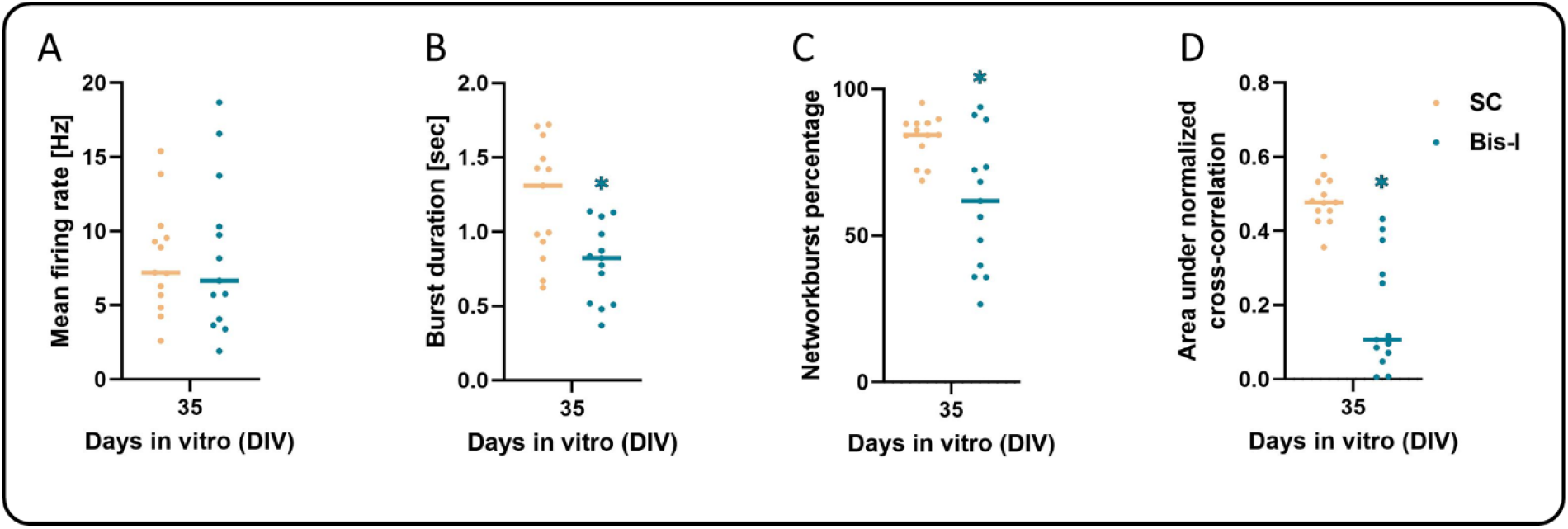
Bisindolylmaleimide I (Bis-I) inhibits neural network development on 48-well microelectrode arrays (MEA) after 35 days of exposure, described by evaluation of specific network parameters. Starting at DIV 7 networks were treated with 5 µM Bis-I and compared to the solvent control (SC) of the respective plate. Data are represented as single experiment values (median of 3 wells each) of 13 independent experiments and merged by median (coloured bar). Statistical significance was calculated using two-tailed Student’s t-tests. A p-value below 0.05 was termed significant. *significant compared to the respective SC.

